# Identification and genetic characterization of MERS-related coronavirus isolated from Nathusius’ pipistrelle (*Pipistrellus nathusii*) near Zvenigorod (Moscow region, Russia)

**DOI:** 10.1101/2022.06.09.495421

**Authors:** A.S. Speranskaya, I.V. Artyushin, A.E. Samoilov, E.V. Korneenko, K. Khabudaev, E.N. Ilina, A.P. Yusefovich, M.V. Safonova, A.S. Dolgova, A.S. Gladkikh, V.G. Dedkov, Daszak Peter

## Abstract

The majority of emerging infectious diseases are caused by pathogens with zoonotic origin, and most of these emerged from wildlife reservoirs. Bats are diverse, and widely distributed globally, and are the known or hypothesized reservoir of a series of emerging zoonotic viruses. Analyses of bat viromes have been used to identify novel viruses with potential to cause human infection. We characterized the fecal virome of 26 samples collected from six bat species captured during 2015 in Moscow Region. Of these 13/26 (50%) samples were found to be coronavirus positive. We sequenced and assembled the complete genome of a novel MERS-related Betacoronavirus from *Pipistrellus nathusii*, named MOW-BatCoV strain 15-22. Of *P. nathusii* 3/6 samples were found to carriers of MOW-BatCoVs. The genome organization of MOW-BatCoV/15-22 was identical to other known MERS-related coronaviruses. Phylogenetic analysis of whole genomes suggests that MOW-BatCoV/15-22 falls into a distinct subclade closely related to human and camel MERS-CoV, and MERS-related CoVs from the bat species *Hypsugo savii* and *Pipistrellus kuhlii* (from Italy) and *Neoromicia capensis* (from South Africa). Unexpectedly, phylogenetic analysis of the novel MOW-BatCoV 15-22 spike genes showed the closest similarity to a bat CoV Neoromicia/5038 and CoVs from *Erinaceus europaeus* (the European hedgehog), thus MOW-BatCoV could arise as result of recombination between ancestral viruses of bats and hedgehogs. Computer molecular docking analysis of MOW-BatCoV 15-22 Spike glycoprotein binding to DPP4 receptors of different mammal species predicted highest binding interaction with DPP4 of the bat *M. brandtii* (docking score -320.15) and the European hedgehog, *E. europaeus* (docking score -294.51). Hedgehogs are widely kept as pets, and are commonly found in areas of human habitation. Our finding of a novel bat-CoV likely able to infect hedgehogs suggests the potential for hedgehogs to act as intermediate hosts for bat-CoVs between bats and humans.

## Introduction

Coronaviruses (CoVs) have been responsible for three high impact outbreaks in the past 2 decades, including severe acute respiratory syndrome (SARS), the Middle East respiratory syndrome (MERS) and the ongoing coronavirus disease 2019 (COVID-19) pandemic. Each of these diseases affects the human respiratory system, causing a spectrum from asymptomatic and mild respiratory illness to severe pneumonia, acute respiratory failure and death.

The current COVID-19 pandemic is now in its third year and continues to be a global health emergency, with more than 500 million confirmed cases, including more than six million deaths, reported to WHO by the date of submitting this manuscript [1]. The earlier outbreak of MERS was caused by zoonotic virus, MERS-CoV, transmitted to humans from infected dromedary camels, with secondary human-to-human transmission mainly nosocomial [2]. MERS was first identified in Saudi Arabia in 2012 [3] and has now been reported in 27 countries, with the largest outbreaks in Saudi Arabia, United Arab Emirates, and the Republic of Korea leading to 858 known deaths due to the infection and related complications The disease has a high fatality rate of up to 35%[4,5]. The origin of the virus is not fully understood yet but phylogenetic analysis of different virus genomes suggest it originated in bats and passed to humans in 2012 after circulating endemically in dromedary camels for around 30 years [6,7].

Bats are the known or putative reservoirs of several viruses that can cause severe human disease, including rabies, Hendra, Nipah, Marburg, SARS, MERS and Ebola viruses [8–10]. Bats also carry diverse coronaviruses, some of which are ableto bind to human cells *in vitro*, suggesting that bats are the likely reservoirs of potential future zoonotic CoVs [11]. In addition, bats are a diverse group of mammals (representing around 1/5th of all mammalian biodiversity), have wide geographic distribution, long life span and are known to feed and roost near to human communities, suggesting they are an important reservoir of other potential emerging diseases [12].

Two genera of CoVs infect mainly mammals: *Alphacoronavirus* (α-CoV) and *Betacoronavirus* (β-CoV) [11,13,14]. Phylogenetic analysis shows that β-CoVs are grouped into several clades (from A to D). These four Betacoronavirus lineages have been reclassified into five subgenera: Embecovirus (A), Sarbecovirus (B), Merbecovirus (C), Nobecovirus (D), and an additional lineage Hibecovirus [15],[16]. MERS CoV, which causes fatal pneumonia in humans and has a dromedary camel reservoir, along with MERS-related (MERSr-) CoVs form clade C (or Merbecoviruses) within the *Betacoronaviruses* and includes CoVsdiscovered in bats and hedgehogs [14–17]. MERSr-CoVs have been reported from South Africa (NeoCoV from *Neoromicia capensis*) [18], Mexico (Mex_CoV-9 from *Nyctinomops laticaudatus*) [19], Uganda (MERSr-CoV PREDICT/PDF-2180 from *Pipistrellus cf. hesperidus*) [20], Netherlands (NL-VM314 from *Pipistrellus pipistrellus*) [21], Italy (BatCoV-Ita1 strain 206645–40 from *Hypsugo savii* and BatCoV-Ita2 strain 206645-63 from *Pipistrellus kuhlii*) [17] and China (BatCoV/SC2013 from *Vespertilio superans* [22] and strains of HKU4- and HKU5-CoVs from *Tylonycteris* and *Pipistrellus* bats [23–26]. To date, few surveys for CoVs have been conducted in Russia, and no MERSr-CoVs have been reported.

Coronavirus cell tropism and ability to infect hosts is determined primarily by the spike protein which is part of the receptor binding domain (RBD) of the sCoV genome [27–29]. Unlike SARS-CoV and SARS-CoV-2, which bind to Angiotensin-converting enzyme 2 (ACE2), MERS-CoV targets the cell surface receptor Dipeptidyl peptidase 4 (DPP4, also known as CD26). DPP4 is relatively conserved among mammalian species, so that MERS-CoV is capable of infecting a wide range of cell lines derived from humans, non-human primates, bats, swine, horse, rabbit, civet, and camel but not from mice, hamster, dog, ferret, and cat [10,27]. The MERS-CoV spike protein undergoes adaptive evolution when inoculated onto normally non-permissive hamster cells transfected to express DPP4 from different bat species [27,30]. HKU4 MERSr-CoVs from bats in China have RBDs that potentially can bind human DPP4 [31] with even low affinity for human cells suggesting potential to infect humans and adapt to more efficient cell entry [17].

In this paper, we describe a survey of bats in Russia for CoVs, report a novel MERSr-CoV, describe its genome organization and relationship with known coronaviruses.

## Materials and methods

### Sample collection

In summer 2015 fecal samples were collected from 26 bats of the following species: *Myotis dasycneme* (n=5), *Myotis daubentonii* (n=5), *Myotis brandtii* (n=3), *Nyctalus noctula* (n=4), *Pipistrellus nathusii* (n=6), *Plecotus auritus* (n=2), *Vespertilio murinus* (n=1), inhabiting the Zvenigorodsky district of the Moscow region (Sharapovskoe forestry, coordinates N55.69, E36.70). No bats were killed for this study and all bats were captured in mist nets and released at the site of capture. Bat capture and sampling was conducted by professionally trained staff of the biological department of Lomonosov Moscow State University. Fecal samples, rectal swabs and ectoparasites were collected after capture, and species, sex, reproductive and health status visually determined by trained field biologists. Swab samples were kept in a transport media for transportation and storage of swabs with mucolytic agent (AmpliSens, Russia) at 4°C during transport to the laboratory, and were then stored at −80 °C before processing.

### RNA extraction and reverse transcription

RNA was extracted from bat faecal samples using QIAamp Viral RNA Mini Kit (Qiagen, Germany). Carrier RNA was dissolved in buffer AVE and added to buffer AVL according to manufacturer’s recommendations before extraction. 140 ul of fecal sample was added to the prepared AVL buffer with carrier RNA–Buffer AVE. Further steps were performed according to the original protocol. RNA was eluted with 60 μL of the AVE Buffer and stored at −70 °C until evaluation. 10 ul of RNA was taken to reverse transcription using Reverta-L (AmpliSens, Russia). Second strand of cDNA was obtained using NEBNext Ultra II Non-Directional RNA Second Strand Module (E6111L, New England Biolabs). In order to increase input concentration 24 ul of first strand product were added to 10 ul of the H2O (milliQ) for the further steps.

### PCR and NGS screening of faecal samples for CoV RNA

PCR-screening for CoV RNA was performed using primers targeting Alpha- and Betacoronavirus species: 5’-CTTATGGGTTGGGATTATCC (CoV2A-F) and 5’-TTATAACAGACAACGCCATCATC (CoV2A-R), described in [32–34]. This generated ∽400-500-bp amplicons from the RNA-dependent RNA polymerase (RdRp) gene. The following thermal cycling parameters were used: 94°C for 3 min, followed by 10 cycles of 94°C for 20 s, 55°C to 45°C (–1°C per cycle) for 20 s, and 72°C for 30 s; and then 42 cycles of 94°C for 20 s, 45°C for 20 s, and 72°C for 30 s; and finally, 72°C for 3 min [35]. PCR amplification products were analyzed by agarose gel electrophoresis. Positive PCR products were purified using AMPure beads and prepared for high throughput sequencing using the TruSeq protocol for Illumina. Sequencing was performed using Illumina MiSeq system to generate 250-bp paired-end reads. Reads were subjected to analysis via the following pipeline: Reads were filtered using Trimmomatic [36]. Then sequences of PCR primers, as well as simple repeats, were masked; filtered reads with an unmasked region of greater than 30 bp were collected and used for taxonomic analysis by comparing assembled contigs or individual reads to the NCBI non-redundant nucleotide and protein sequence databases using the blastn, blastx or tblastn algorithms as described in [35].

### Library preparation and high throughput sequencing of viral genome

Double stranded cDNA was taken to library preparation using NEBNext Ultra II DNA Library Prep Kit for Illumina (New England Biolabs, England). End prep was performed according to the manufacturer’s protocol. For the adaptor ligation step we chose Y-shaped adaptors compatible with Nextera XT Index Kit in the amount of 4 pM per reaction. PCR amplification with index adaptors in the amount of 7.5 pM per reaction was performed with Nextera XT Index Kit (Illumina) in 25 μL total volume according to NEBNext Ultra II DNA Library Prep Kit for Illumina protocol with 15 cycles.

High throughput sequencing was performed using Illumina HiSeq 1500 with HiSeq PE Rapid Cluster Kit v2 and HiSeq Rapid SBS Kit v2 (500 cycles). Paired reads were filtered with Trimmomatic using parameters SLIDINGWINDOW:4:25 MINLEN:40. After read trimming, genome assembly and selection of Coronaviridae sequences, we obtained two contigs with lengths of 20098 and 10135 bp. Genome assembly was completed by SPAdes 3.15.0 [37]. Coronaviridae sequences were selected by BLASTn [38] of assembled contigs using all of the available Coronaviridae genomes as a reference. Read mapping was performed using bowtie2 [39].

Alignment of contigs to the closest MERSr-CoV genomes (MG596802.1) revealed an uncovered 62 bp fragment between these contigs. Gaps within the assembled genome were closed and confirmed using Sanger sequencing. We performed Sanger sequencing of this area to connect two contigs and obtain full-genome sequencing with following primers: 1-forward ACATACGTGACAATGGTTCATTAG, and 1-reverse CTGTTGACTCTCTATAAATATAGAAC.

Genome annotation was performed by Geneious 7.1.9 and edited manually.

TRS-L and TRS-B aligment was made by Vector NTI software

### Phylogenetic analysis

We downloaded beta-CoV sequences from GenBank using the keyword “Merbecovirus” as the primary filter to identify beta-CoVs from bats, humans, camels and other mammals. We included 9 full genomes of CoVs from bats, 1 from a lama, 232 from camels and 254 from humans. For phylogenetic tree construction we used only complete genomes of viruses from highly represented hosts: individual trees were previously constructed separately for full genomes of camel and human Merbecoviruses. Clusters with a p-distance >0,001 were collapsed and one sequence per cluster was selected randomly. The complete genome of the newly discovered CoV was aligned with all sequences using MAFFT v7 via the online service of RIMD (Research Institute for Microbial Disease of Osaka University, Japan): https://mafft.cbrc.jp/alignment/software then bp on 5’-end and 3’-end were trimmed.

For phylogenetic analysis of genes encoding Nucleocapsid protein the partial sequences designated in metadata as “merbecovirus nucleocapsid gene” were downloaded from GenBank and combined with sequences extracted from complete genomes (sampling described above). The obtained 128 sequences were aligned using MAFFT v7, trimmed the ∼110 bp on 5’-end and ∼200 bp on 3’-end. At last, the phylogenetic tree was constructed on alignment of ∼1308 bp, the Best-fit model according to BIC: TIM2+F+I+G4.

For phylogenetic analysis of RdRp-encoding region of ORF/b as well as Spike-genes the similar method of sampling was used, namely: the partial sequences designated in metadata as “merbecovirus RdRp” or “merbecovirus Spike gene” were downloaded from GenBank, (2) combined with relevant sequences extracted from complete genomes, aligned using MAFFT v7, trimmed in the 5’-end and 3’-end. For the RdRp-encoding region, the ∼2801 bp alignment of 118 sequences was used for phylogenetic tree construction, the Best-fit model according to BIC: GTR+F+I+G4. For the Spike, the ∼4026 bp alignment of 123 sequences was used for phylogenetic tree construction, the Best-fit model according to BIC: GTR+F+I+G4.

Phylogenetic analyses were performed using W-IQ-TREE (http://iqtree.cibiv.univie.ac.at/) with ModelFinder [40], tree reconstruction [41], and ultrafast bootstrap (1000 replicates) [42]. Phylogenetic trees and coevolutionary events were visualized using the online website (https://itol.embl.de/) with iTOL software [43].

### Structural modeling and molecular docking

The SWISS-MODEL server [44] was used to determined the three-dimensional structure of the MOW-BatCoV spike protein, and the structures of DPP4 for *Myotis brandtii* [EPQ03439.1], *Pipistrellus kuhlii* (KAF6353216.1), *Erinaceus europaeus* [XP_016043930.1], *Felis catus* [NP_001009838.1] and *Mus musculus* [NP_034204.1]. The DPP4 sequence of *E. europaeus* contains X in the sequence, which we replaced with the W sequence. The 5T4E crystal structure of DPP4 of Homo sapiens was obtained from the RCSB.

The constructed models and the crystal structure of DPP4s of the studied organisms were docked to the modeled structure of the MOW-BatCoV Spike protein using the HDOCK server [45]. The RBD (360-610 a.r.) of MOW-BatCoV and all DPP4 were used as docking sites. The docking results were analyzed in the PyMOL Molecular Graphics System, Version 2.0 [46].

## Results

### General description

We collected and analyzed 26 fecal samples from six species of bats: *Myotis dasycneme, Myotis daubentonii, Myotis brandtii, Nyctalus noctula, Pipistrellus nathusii, Plecotus auritus*. All bats were visually healthy. Ectoparasite analysis yielded mites in 21/26 samples with 2 samples containing both mites and fleas. PCR fragments of RdRp yielded results in 13 of 26 fecal samples, giving an overall detection rate of 50%. Using Illumina MiSeq high throughput sequencing analysis and data analysis as described in [35], we confirmed presence of CoVs in 8/13 samples (the five remaining smaples were not sequenced). Five of the six species of bats sampled were infected by different CoV strains (*M. dasycneme, M. brandtii, M. daubentonii, N. noctula* and *P. nathusii*), and only one sample was negative for CoVs (from *P. auritus*). Of investigated species the only *Pipistrellus nathusii* were carriers of betacoronaviruses: sample numbers 16, 22 and 33, see table 1. The RdRp fragments were sequenced, all they shared >98% sequence identity to KC243390.1 (BtCoV/8-724/Pip_pyg/ROU/2009) which was previously found in *Pipistrellus pygmaeus* from Romania and published in 2013 [47]. We propose that these represent a potentially novel Betacoronavirus in the *P. nathusii* and sequenced the complete genome of virus from sample №22.

**Table 1.** *P. nathusii* in which natural coronavirus infection has been found. Samples where Betacoronaviruses were detected are marked in bold.

| Sample ID | Gender | Age | Ectoparasites | PCR product | Characterisation of PCR product by NGS |  |  |  |
| --- | --- | --- | --- | --- | --- | --- | --- | --- |
|  |  |  |  |  | fragment length, bp | Genbank ID, nearest | Identity, % | Genus |
| Bat№16 | F | Semi-adult | Mites | + | 418 | **KC243390.1** | **98.35** | **Beta** |
|  |  |  |  |  | 420 | EU375869.1 | 99.51 | Alpha |
| Bat№21 | F | Semi-adult | Mites | + | 420 | EU375864.1 | 98.77 | Alpha |
| **Bat№22** | **M** | **Semi-adult** | **Mites** | **+** | **418** | **KC243390.1** | **98.51** | **Beta** |
| Bat№23 | M | Adult | Mites | + | 420 | EU375869.1 | 99.51 | Alpha |
| Bat№25 | F | Adult | A lot of Mites | - | - | - | - | - |
| **Bat№33** | **M** | **Semi-adult** | **A lot of Mites** | **+** | 420 | EU375864.1 | 98.77 | Alpha |
|  |  |  |  |  | **418** | **KC243390.1** | **98.35** | **Beta** |

*Pipistr*e*llus nathusii* is widely distributed across Europe, see Figure 1 (b). In Figure 1 (a) shows the bat caught in the process of work on the fecal samples collection. The metagenome sequencing of total RNA extracted from Sample №22 resulted in 248.8 million paired reads (SRR15508267), 0.01% of them were mapped to initially obtained Coronaviridae contigs resulting in the complete genome. It was named as MOW-BatCoV strain 15-22 and has been deposited in the GenBank under the accession numbers ON325306. RdRp fragments of viral genomes from two other samples (№16 and №33) have been deposited in the GenBank as Bat-CoV/P.nathusii/Russia/MOW15-16/1/15 (accession number ON676527) and Bat-CoV/P.nathusii/Russia/MOW15-33/1/15 (accession number ON676528).

**Figure 1.**
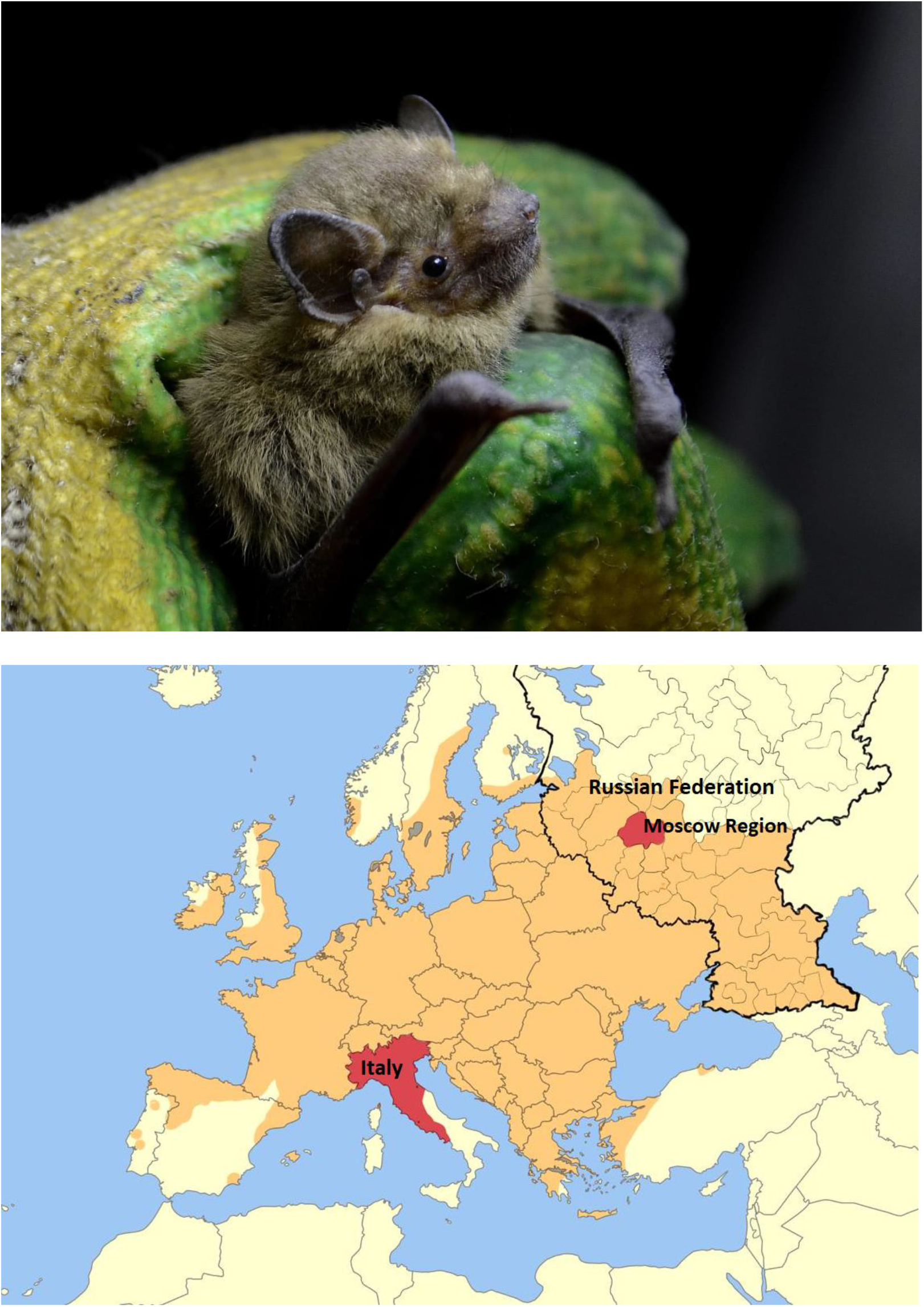
(a) *Pipistrellus nathusii*, also known as Nathusius’ pipistrelle, studied here; (b) Geographical distribution of *Pipistrellus nathusii* in Europe. The dark-yellow labeled region represents their habitats according to [48] and [49]. The red labeled area represents the location where MERS-related Bat-CoV were obtained.

Bat-CoV/P. nathusii/Russia/MOW-BatCoV 15-22/2015 contains 30257 bases, with G+C content 43,72%=, and a genome organization similar to other members of *Merbecovirus*, namely: ORF1ab encoding putative mature nonstructural proteins, including RdRp (RNA-dependent RNA polymerase) — S (spike protein) — the genes encoding nonstructural proteins NSP3, NSP4a, NSP4b and NSP5 — E (envelope protein) — M (membrane glycoprotein) — N (nucleocapsid phosphoprotein) — the gene encoding nonstructural protein NSP8 (Figure 2). Predicted functional domains in the ORFs are summarized in Table 2.

**Table 2.** Genome localization of predicted protein sequences of MOW-BatCoV 15-22 MERS-related coronavirus.

| **ORF** | **nt position (start-end)** | **No. of amino acids** | **putative leader TRS-L and TRS-B** |
| --- | --- | --- | --- |
| ORFa | 175-13524 | 4449 | TRS-L |
| ORFab | 175-21587 | 7137 | TRS-L |
| S prot | 21529-25629 | 1367 | TRS-B |
| ORF3 | 25645-25959 | 105 | TRS-B |
| ORF4a | 25968-26252 | 95 | TRS-B |
| ORF4b | 26173-26925 | 251 | No |
| ORF 5 | 26935-27612 | 226 | TRS-B |
| E prot | 27691-27939 | 83 | TRS-B |
| M prot | 27954-28619 | 222 | TRS-B |
| Nprot | 28675-29982 | 436 | TRS-B |
| ORF 8b | 28721-29314 | 198 | no |

**Figure 2.**
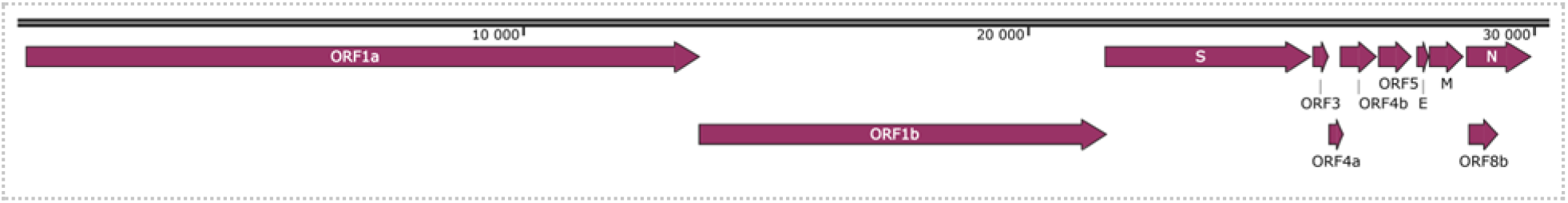
Genome organization of MW-BatCoV, strain 15-22. Nonstructural genes (ORF1ab, ORF3, ORF4a, ORF4b, ORF5, orf8b) and structural genes (S, E, M, N) are illustrated. The major open reading frames (ORFs) had the order: ORF1ab (putative mature nonstructural proteins, including RdRp (RNA-dependent RNA polymerase)) — S (Spike) — the genes encoding nonstructural proteins (NS3, NS4a, NS4b, NS5) — E (envelope) — M (membrane glycoprotein) — N (nucleocapsid phosphoprotein) — the gene encoding nonstructural protein NS8b.

The size and genomic localization of the nonstructural proteins (NSP 1–16) encoded by ORF1ab were predicted by sequence comparison with MERS CoV (human HCoV-EMC/2012) and other beta-CoV species. Proteins and 15 expected cleavage sites are shown in Table 3. In ORF1ab the sequence “UUUAAAC”, which is conserved throughout all CoVs -is located at nucleotide position 13497–13503. A predicted leader transcription regulatory sequence (TRS-L), and seven putative transcription regulatory sequences body TRS-B, representing signals for the discontinuous transcription of subgenomic mRNAs (sgmRNAs), have been identified (Table 3). All TRS have conserved AACGAA motif forming the conserved TRS core in betacoronaviruses [50]. With only one G/A modification in ORF3 (Figure 3).

**Table 3.**
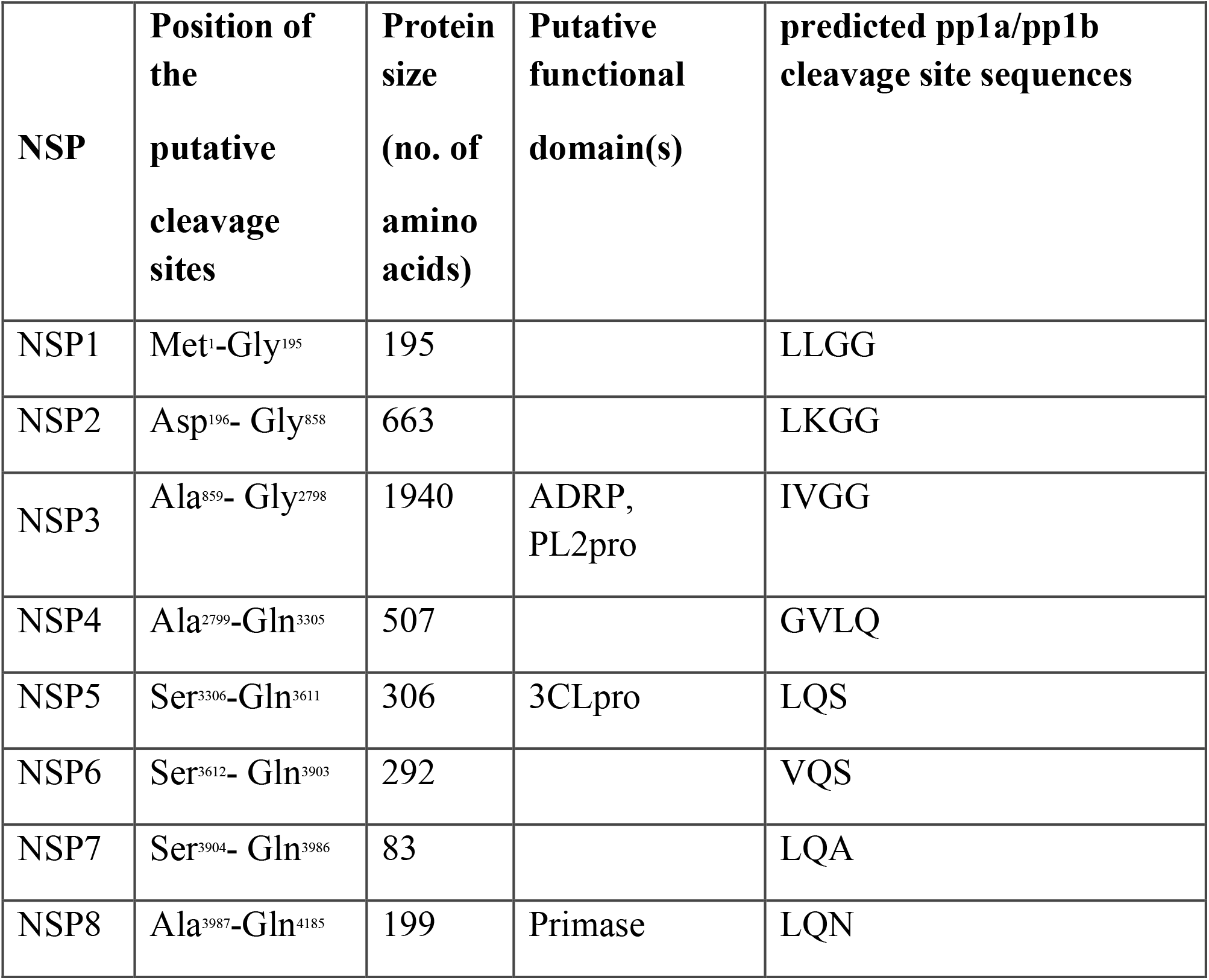
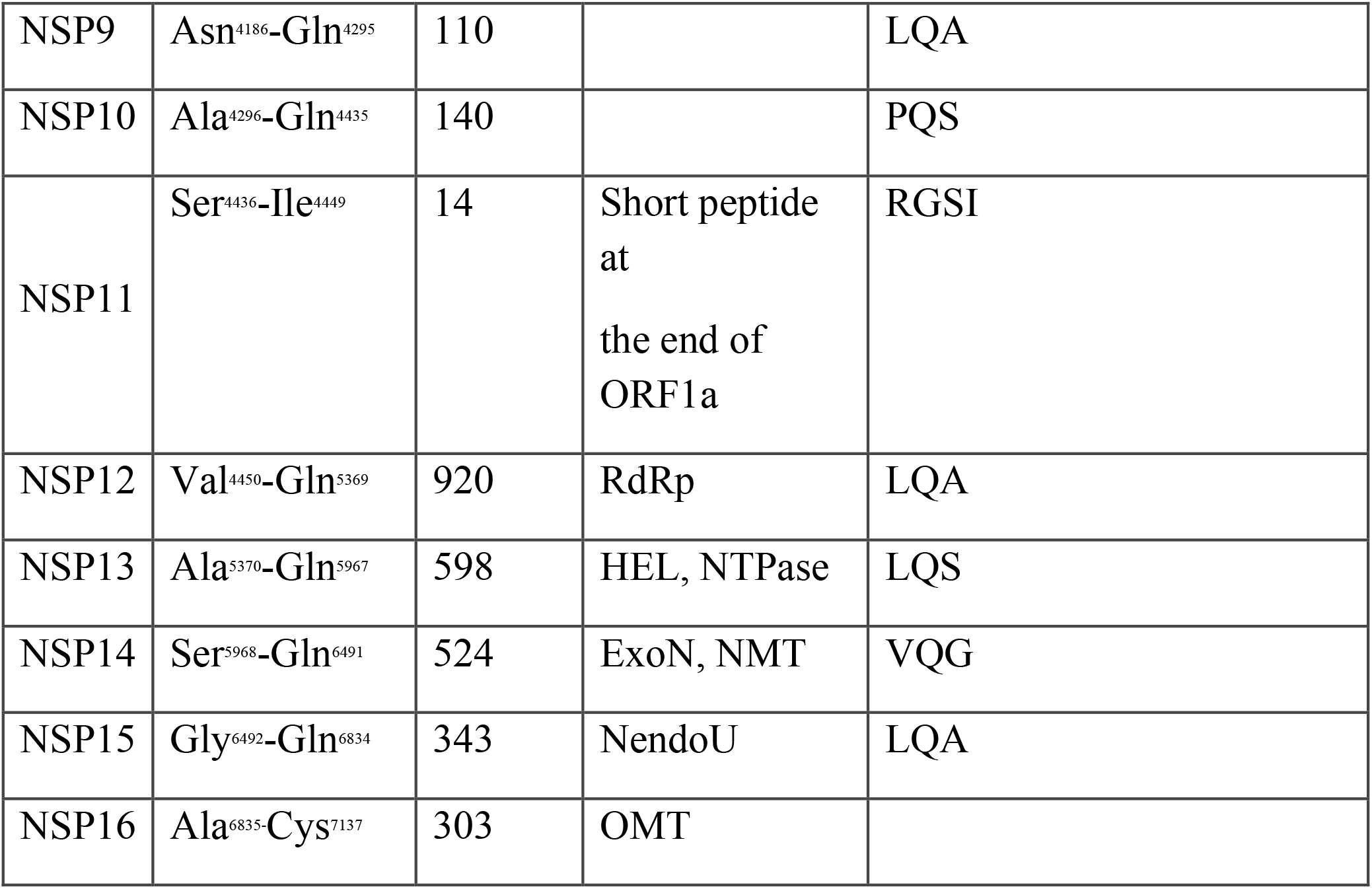
The list of proteins and 15 expected cleavage sites encoded by ORF1ab of MOW-BatCoV 15-22 MERS-related coronavirus. Superscript numbers indicate positions in polyprotein pp1a/pp1ab or position in available sequence with the supposition of a ribosomal frameshift based on the conserved slippery sequence (UUUAAAC) of Coronaviruses. Localized at nucleotide position 13497–13503 ADRP - ADP-ribose 1-phosphatase, PL2pro - papain-like protease 2, 3CLpro - coronavirus NSP5 protease, Hel - helicase, NTPase - nucleoside triphosphatase, ExoN – exoribonuclease, NMT N7 - methyltransferase, NendoU - endoribonuclease, OMT - 2’ O-methyltransferase.

**Figure 3.**
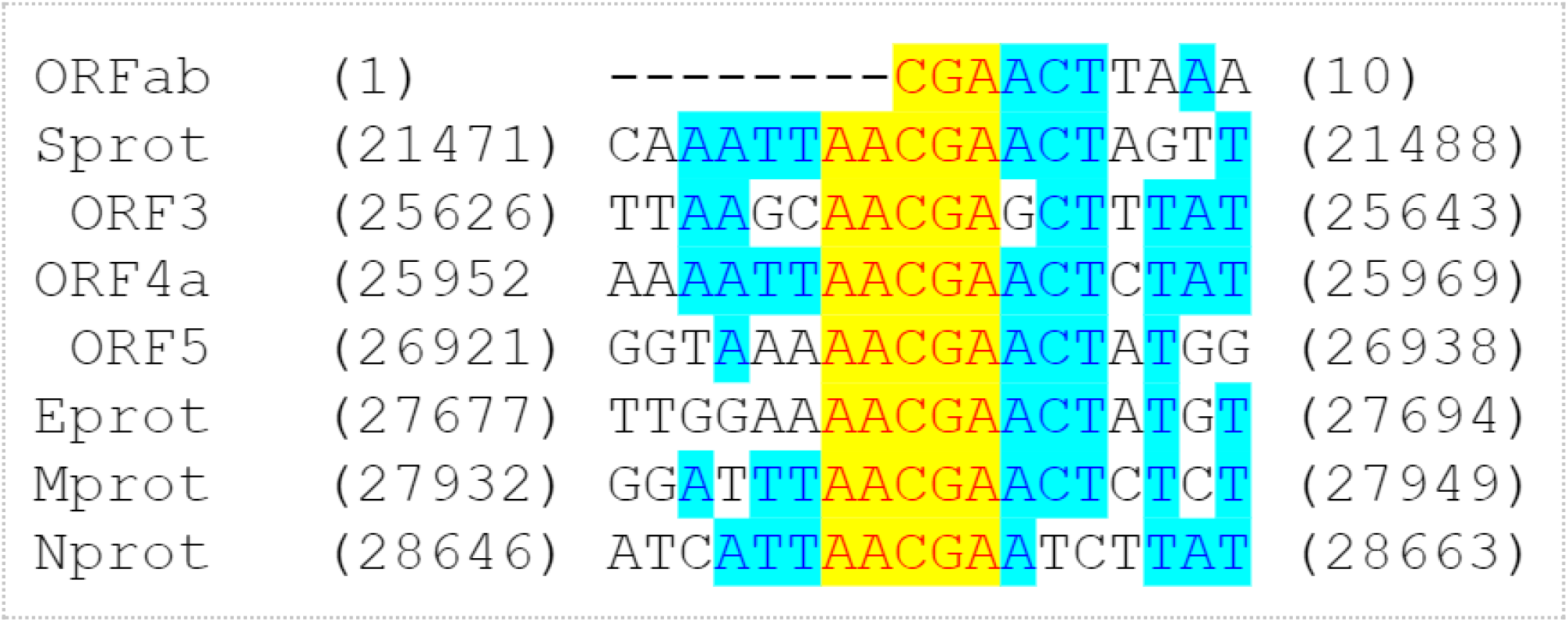
Putative leader TRS-L and TRS-B alignment. Strictly conservative and identical residues are highlighted in yellow and blue, respectively.

The ICTV proposed that viruses sharing >90% aa sequence identity in the conserved replicase domains should be considered conspecific [51]. A separate comparison of the amino acid sequences of seven conserved ORF1ab domains, as suggested by the ICTV) for formal CoV species delineation was made and only the NSP3 (ADRP) aa sequence is below the 90% threshold value in comparison with MERS. The ORF1ab possessed 81,5-82,3% na identities to the ORF1ab of other members of *Merbecovirus*. Comparison of the seven conserved domains in replicase polyprotein pp1ab with other coronaviruses showed that MOW-BatCoV 15-22 possessed for NSP3 (ADRP) - 68,7% of homology, NSP5 (3CLpro) - 90,8%, NSP12 (RdRp) - 96,6%, NSP13 (Hel) - 97,8%, NSP14 (ExoN) - 97,9%, NSP15 (NendoU) - 93,3% and NSP16 (O-MT) - 93,7% aa identities to other members of *Merbecovirus* respectively.

MOW-BatCoV 15-22 MERSr-CoV has nucleic acid identity from 81,32% to 82,46% (with coverage 82-85%) to the ten closest MERS or MERSr-CoVs from bats (*Vespertilio sinensis, Vespertilio superans, Pipistrellus cf. hesperidus*) humans and camels reported between 2013 and 2015 (Table 4). The highest sequence identity was to 2 MERSr-CoVs from bats (*Hypsugo savii* and *Pipistrellus kuhlii*) captured in Italy in 2011 (88% coverage genomes were 81.48% and of 81.45%, respectively).

**Table 4.** Analysis of full genome identify among MOW-BatCoV 15-22 and other members of Merbecovirus (the top 10 of closest full genome sequences available in GenBank).

| Description | Given Scientific Name (by Author's) | Host | Collection date | Country | Query Cover | Per. ident | Acc. Len | Accession |
| --- | --- | --- | --- | --- | --- | --- | --- | --- |
| Bat-CoV/H.savii/Italy/206645-40/2011, complete genome | Middle East respiratory syndrome-related coronavirus | *Hypsugo savii* (bat) | 2011 | Italy | 88% | 81,48% | 30048 | MG596802.1 |
| Bat-CoV/P.khulii/Italy/206645-63/2011, complete genome | Middle East respiratory syndrome-related coronavirus | *Pipistrellus kuhlii* (bat) | 2011 | Italy | 88% | 81,45% | 30039 | MG596803.1 |
| Bat coronavirus Vs-CoV-1 genomic RNA, nearly complete genome | Bat coronavirus | *Vespertilio sinensis* (bat) | undescribed | Japan: Tokyo, Okutama | 85% | 81,32% | 29930 | LC469308.1 |
| BtVs-BetaCoV/SC2013, complete genome | BtVs-BetaCoV/SC 2013 | *Vespertilio superans* (bat) | 2013 | China | 82% | 81,27% | 30423 | KJ473821.1 |
| isolate PREDICT/PDF-2180, complete genome | Bat coronavirus | *Pipistrellus cf. hesperidus* (bat) | 2013 | Uganda | 82% | 80,84% | 29642 | NC_034440.1 |
| isolate Riyadh_9_2013, complete genome | Middle East respiratory syndrome-related coronavirus | *Homo sapiens* | 2013 | Saudi Arabia | 82% | 82,47% | 30055 | KJ156869.1 |
| isolate Hu/Jordan-201440011123/2014, complete genome | Middle East respiratory syndrome-related coronavirus | *Homo sapiens* | 2014 | Jordan | 82% | 82,46% | 30123 | MK039552.1 |
| strain<br>Hu/UAE_032_2014<br>, complete genome | Middle East<br>respiratory<br>syndrome-related<br>coronavirus | *Homo*<br>*sapiens* | 2014 | United<br>Arab<br>Emirate | 82% | 82,46% | 30123 | KY581693.1 |
| camel/UAE_B73_2<br>015, complete<br>genome | Middle East<br>respiratory<br>syndrome-related<br>coronavirus | *Camelus*<br>*dromedarius* | 2015 | United<br>Arab<br>Emirates | 82% | 82,46% | 30123 | MF598663.1 |
| isolate<br>Florida/USA-<br>2_Saudi<br>Arabia_2014,<br>complete genome | Middle East<br>respiratory<br>syndrome-related<br>coronavirus | *Homo*<br>*sapiens* | 2014 | USA | 82% | 82,46% | 30123 | KP223131.1 |

### Phylogenetic analysis

Our phylogenetic analysis of complete MERSr-CoV genomes showed four distinct clades (Figure 4). Clade I consist of nine CoV sequences from hedgehogs. Clade II consists of 17 CoV sequences from bats. Clade III consists of 59 sequences belonging to two subclades: (a) 9 CoV sequences from bats; (b) 49 CoV sequences from humans and camels, as well as one from a bat (strain Neoromicia/5038, from virome of the bat *Neoromicia capensis* collected in South Africa, in April of 2015, Genbank accession number MF593268.1). MOW-BatCoV 15-22 belonged to Clade III(a) along with MERSr-CoVs from *H. savii* and *P. kuhlii* from Italy.

**Figure 4.**
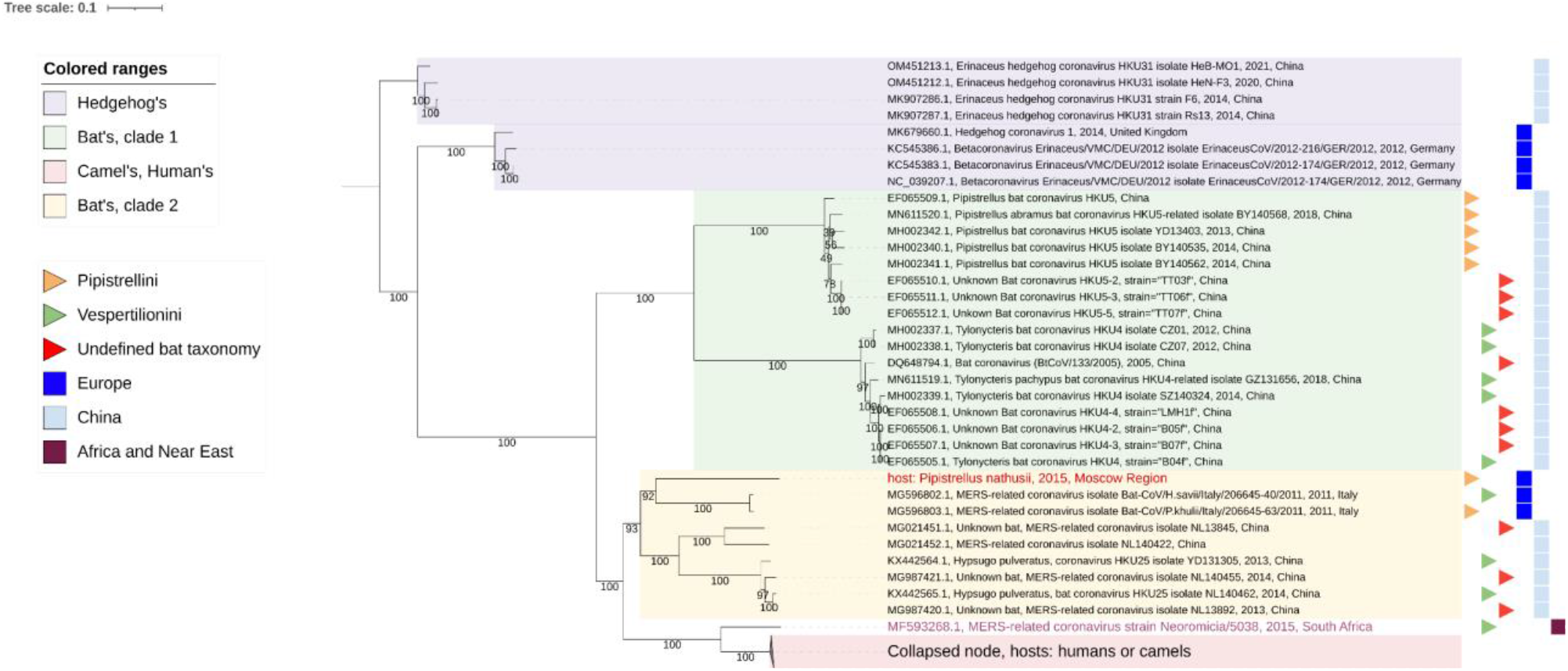
Phylogenetic trees of the Merbecovirus genomes (from complete genomes only). The phylogenetic tree was constructed on 84 complete genome sequences, excluding 5’- and 3’-ends (29757-30331 bp). Numbers show bootstrap values. Best-fit model of substitution according to BIC: GTR+F+I+G4. The virus described in this study is labeled in red bold font.

The phylogenetic trees of Nucleocapsid (N) and RdRp encoding sequences (Figures 5a, 5b) show that clade I consists of CoVs from hedgehogs, clade II consists of the bat CoVs from China, represented by two separate subclades described in [25]. Clade III is represented by two separate subclades: (a) 6 CoVs from bats and (b) human and camel MERS-CoV and a bat CoV. MOW-BatCoV 15-22 belongs to Clade III(b), consisting of numerous human and camel MERS sequences (branches are collapsed in Figure 5) and a small number of bat-CoVs. In the N-tree the MOW-BatCoV 15-22 forms a distinct subclade closely related to human and camel MERS-CoV and MERSr-CoVs from *H. savii* and *P. kuhlii* (from Italy) and MERSr-CoV strain Neoromicia/5038 obtained from *Neoromicia capensis* (Common names: Cape Serotine Bat, inhabits South Africa) (see Figure 5a).

**Figure 5a.**
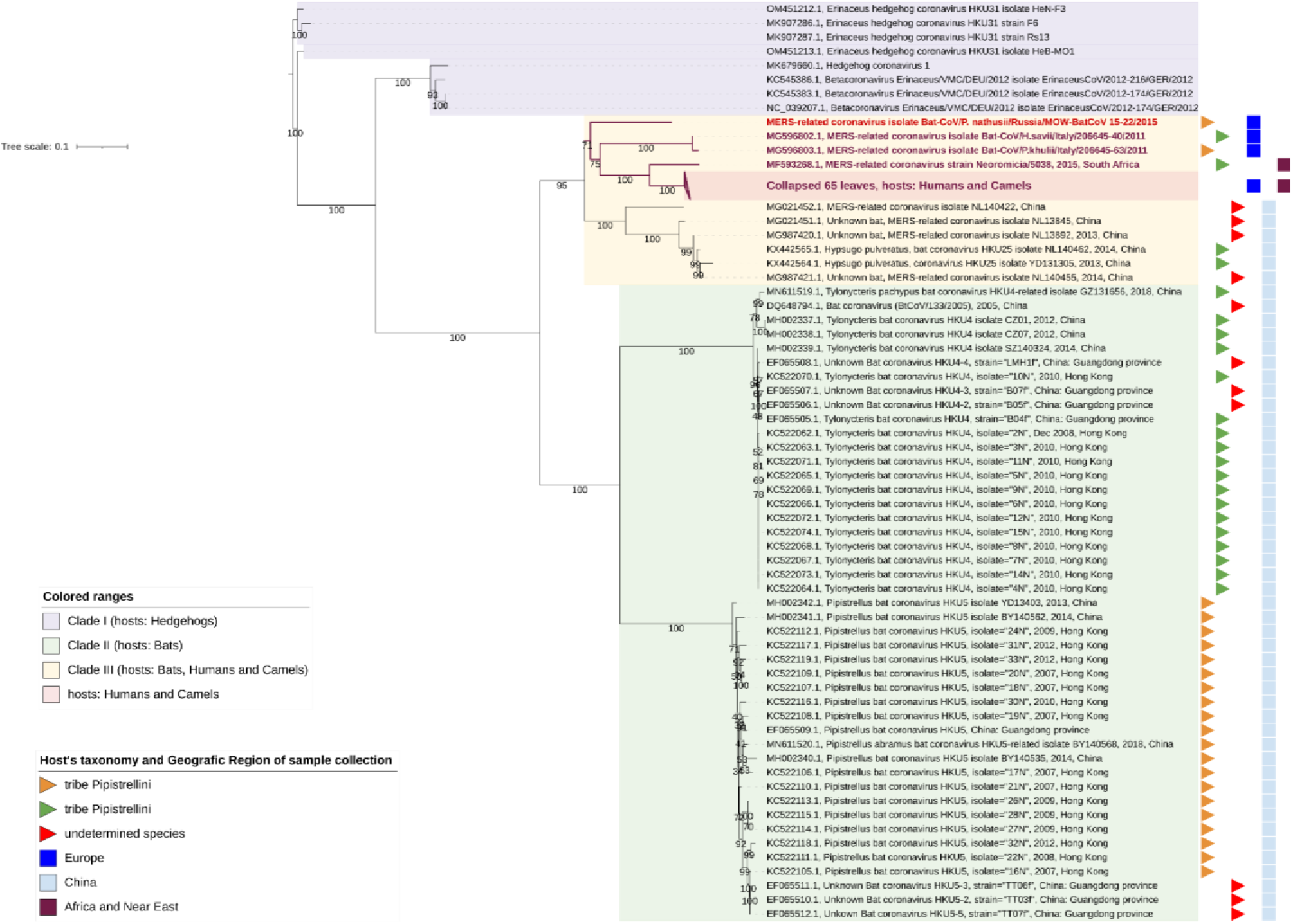
Phylogenetic trees of the Nucleocapsid protein coding regions of Merbecovirus genomes. The phylogenetic tree was constructed on 65 partial N-gene sequences (1272-1297 bp). Numbers show bootstrap values. Best-fit model of substitution according to BIC: TIM2+F+I+G4. The virus described in this study is labeled in red bold font.

**Figure 5b.**
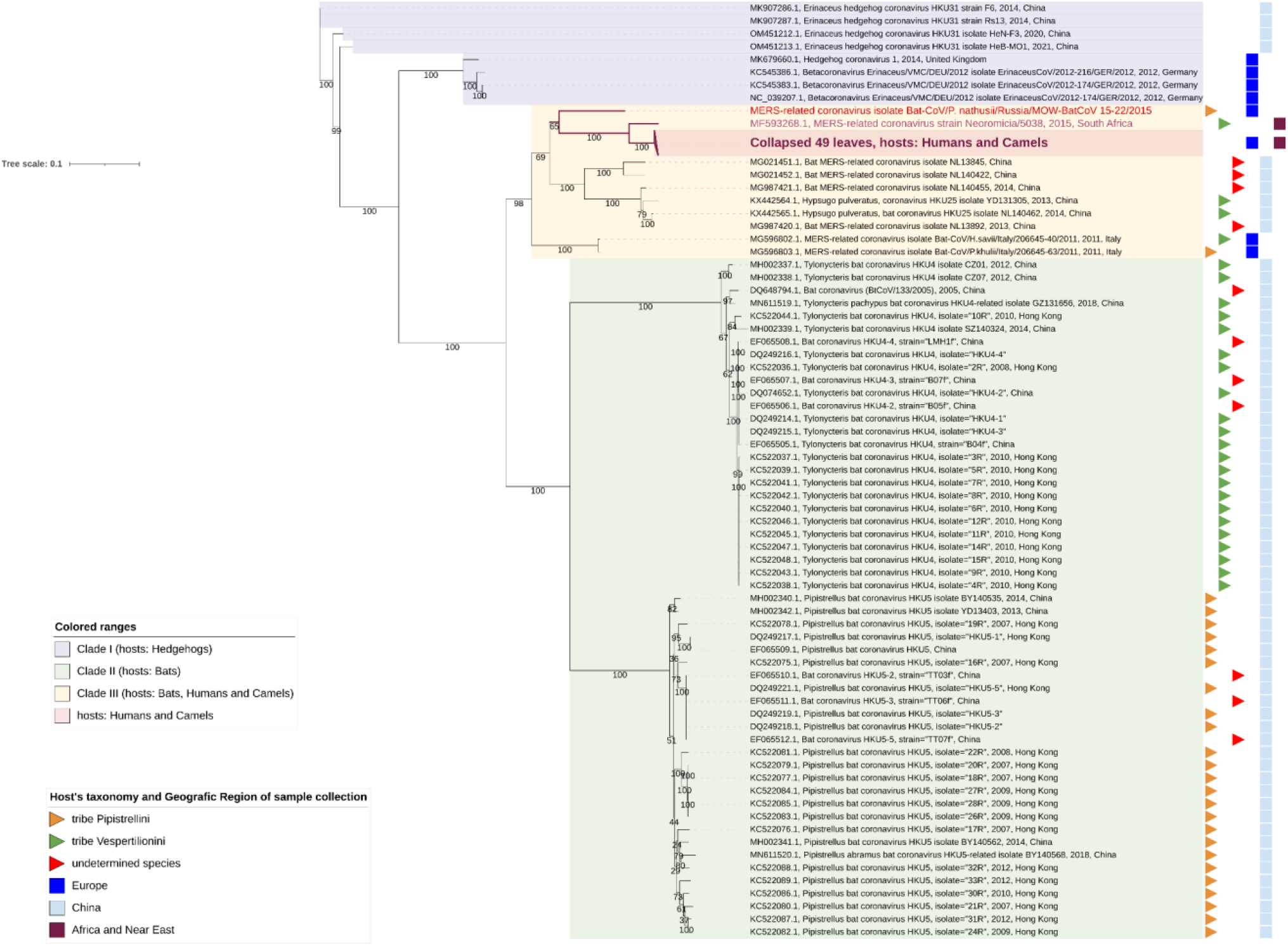
Phylogenetic tree based on 2798-2802 of the partial RdRp (NSP12) proteins coding regions of Merbecovirus genomes. Numbers show bootstrap values. Best-fit model of substitution according to BIC: GTR+F+I+G4. The virus described in this study is labeled in red bold font.

**Figure 5c.**
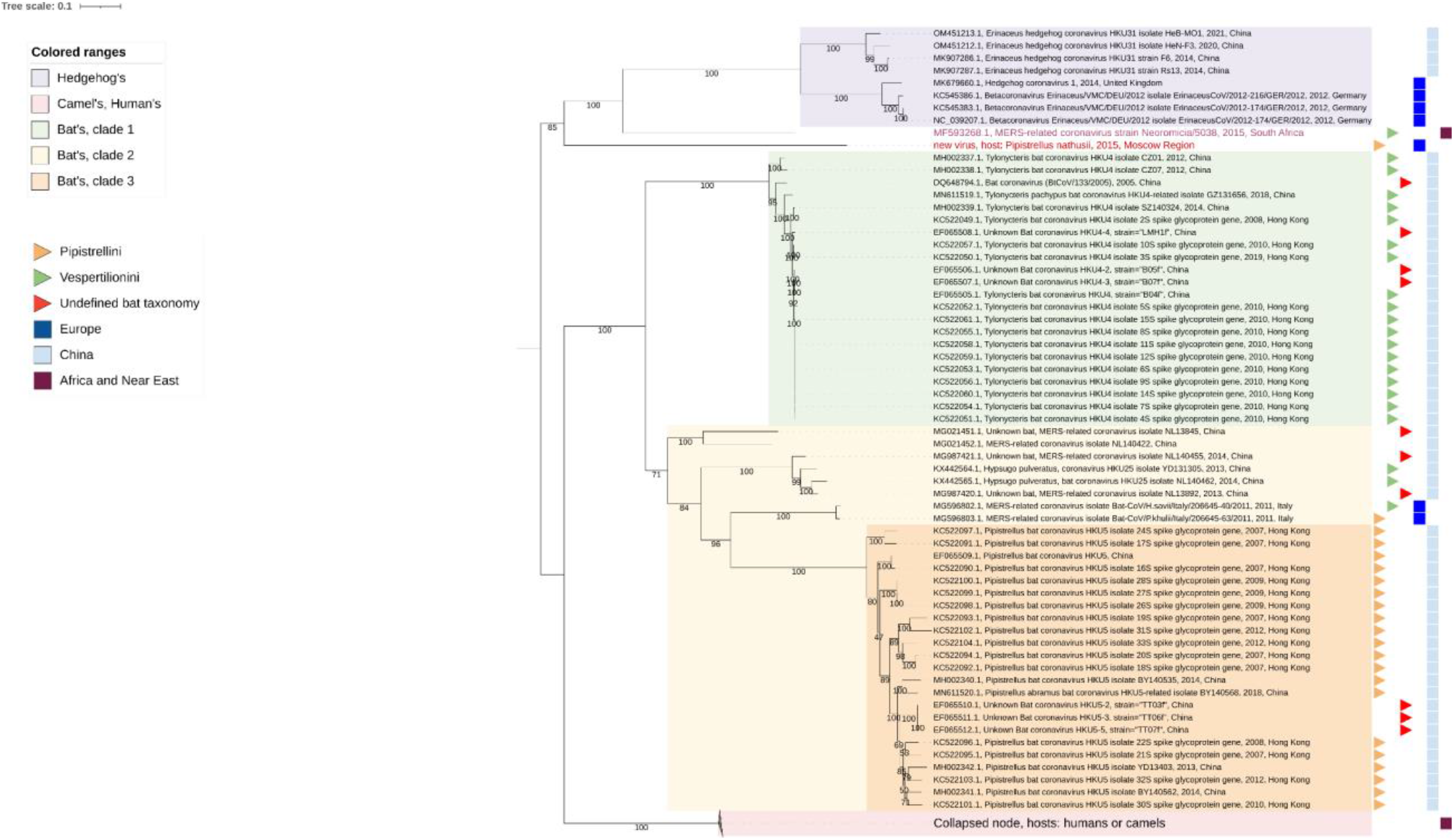
Phylogenetic tree based on the partial Spike glycoprotein protein encoding regions of Merbecovirus genomes. Numbers show bootstrap values. Best-fit model of substitution according to BIC: GTR+F+I+G4. The virus described in this study is labeled in red bold font.

In the RdRp-tree the MOW-BatCoV 15-22 formed a subclade MOW-BatCoV 15-22 together with the only Neoromicia/5038 and human/camel’s viruses, but MERS-related coronaviruses from *H. savii* and *P. kuhlii* (from Italy) fall into different clade (see Figure 5b).

Phylogenetic analysis of the spike protein encoding region of Merbecoviruses (Figure 5c) shows that MOW-BatCoV 15-22 is most closely related to coronavirus strain Neoromicia/5038. Unexpectedly, both MOW-BatCoV 15-22 and Neoromicia/5038 showed closest S-protein sequence identity to CoVs from *Erinaceus* hedgehogs, forming a distinct branch among Merbecovirus (Figure 5c).

### Docking

To predict and analyze the interaction of MOW-BatCoV Spike glycoprotein with DPP4 receptors of the different mammalian species, the three-dimensional structures of these proteins were obtained by homologous modeling. The DPP4 proteins of two bats (*Myotis brandtii* and *Pipistrellus kuhlii*), the hedgehog (*Erinaceus europaeus*), domestic cat (*Felix catus*) and mouse (*Mus musculus*) were used for analysis.

MOW-BatCoV Spike protein structure was determined using the 6Q04 reference structure of human MERS-CoV Spike protein. We selected the protein structure of the amino acid sequence with the highest homology (61.41%) to the input sequence as a template. The resulting model had a GMQE of 0.61.

For the DPP4 structure models of mammals, the reference human FAPalpha was used. The best 6Y0F structure models were determined for *M. brandtii* and *P. kuhlii* DPP4 with 92.14% identity (GMQE = 0.91) and 92.31% identity (GMQE = 0.93), respectively. The best reference structure for DPP4 structure models of *E. europaeus, F. catus*, and *M. musculus* was 2QT9 with identities of 84.97% (GMQE = 0.9), 87.97% (GMQE = 0.89) and 84.15% (GMQE = 0.9), respectively.

Molecular docking allowed us to determine the best binding cluster for the Spike-protein of MOW-BatCoV and DPP4 proteins of the organisms studied. Highest binding was predicted between MOW-BatCoV spike protein and DPP4 of *M. brandtii* (docking score -320.15). The second highest binding was predicted for *E. europaeus* (docking score -294.51), which is consistent with phylogenetic analyses. Docking results predicted higher binding of the MOW-BatCoV S to *H. sapiens* DPP4 (docking score -290.79) than to *P. kuhlii* DPP4 (−274.21), *M. musculus* (docking score -262.74) and *F. catus* (docking score -248.18), see Figure 6 (a-f).

**Figure 6.**
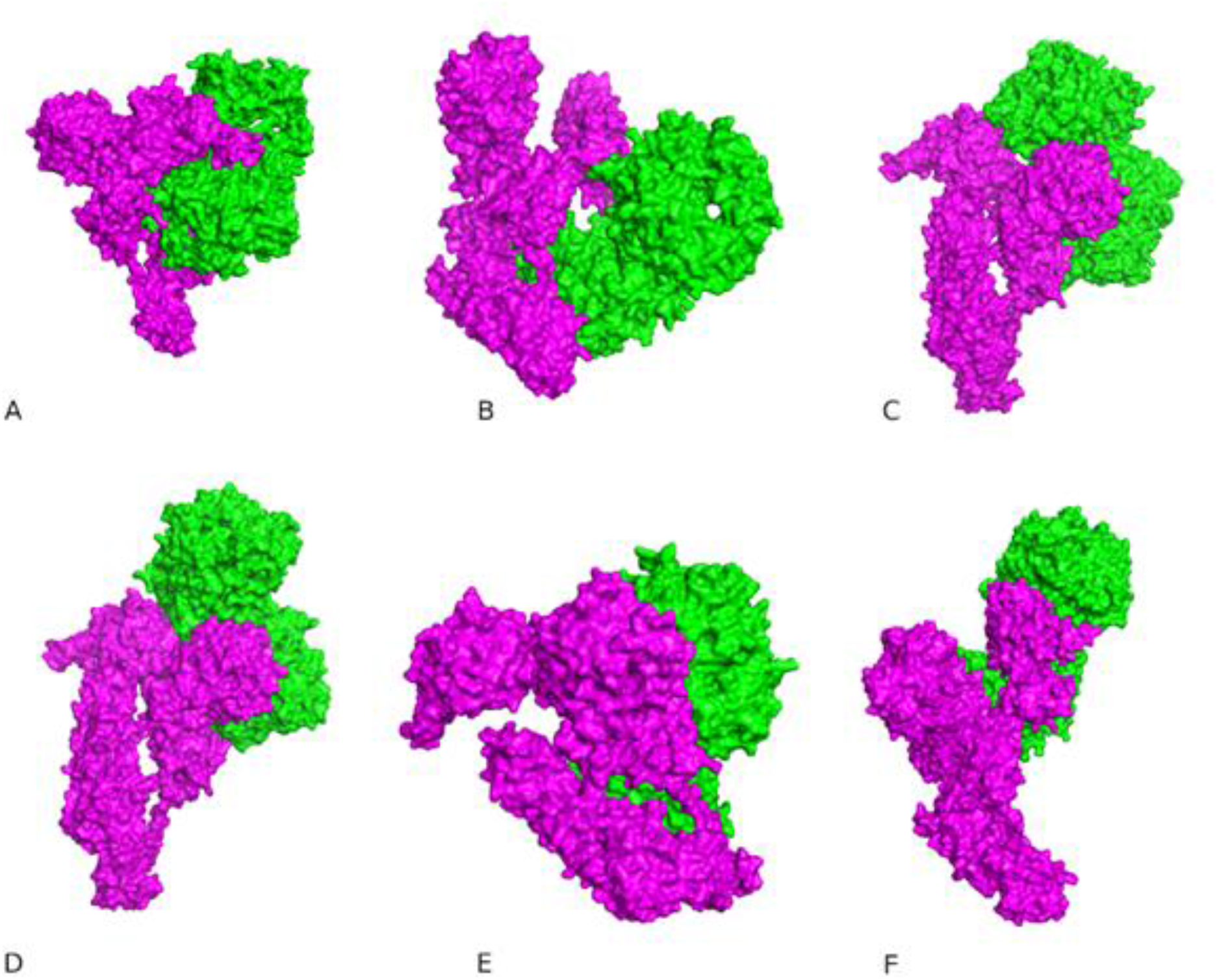
Protein–protein binding interactions of MOW-BatCoV Spike protein and DPP4: A - *M. brandtii*, B - *P. kuhlii*, C - *E. europaeus*, D - *F. catus*, E -*M. musculus*, F - *H. sapiens*. The MOW-BatCoV Spike proteins are marked in magenta, DPP4 are marked in green.

Analysis of the results of molecular docking of the Spike-protein of MOW-BatCoV and DPP4 protein of *M. brandtii* and *E. europaeus* showed that the Spike-protein of MOW-BatCoV shares binding sites with DPP4 of these two species. Thus, 43 and 42 binding sites were shown for *M. brandtii* and *E. europaeus* in the Spike-protein of MOW-BatCoV, among which 31 sites were identical (Table 5).

**Table 5.** Predicted MOW-BatCoV-DPP4 binding sites of *M. brandtii* and *E. europaeus*.

|  | *Myotis brandtii* | *Erinaceus europaeus* |
| --- | --- | --- |
| binding sites | 36, 163, 165, 166, 188, 189, 191, 230, 232, 233, 370, 389, 417, 419, 485, 486, 487, 488, 489, 490, 585, 611, 613, 617, 618, 619, 620, 637, 696, 697, 698, 701, 703, 704, 705, 707, 716, 718, 719, 720, 722, 724, 734 | 36, 38, 39, 154, 156, 157, 158, 159, 163, 165, 166, 188, 189, 190, 191, 197, 230, 232, 233, 370, 389, 416, 417, 419, 485, 486, 487, 488, 489, 490, 585, 613, 617, 618, 619, 620, 697, 698, 699, 703, 704, 718 |
| overlapping binding sites | 36, 163, 165, 166, 188, 189, 191, 230, 232, 233, 370, 389, 417, 419, 485, 486, 487, 488, 489, 490, 585, 613, 617, 618, 619, 620, 697, 698, 703, 704, 718 |  |

## Discussion

A large number of β-CoVs has been identified from bats globally. The MERSr-CoV described here is the first to be identified in Russia. Twenty-six animals of six different bat species which wide distributed in the central European part of the Russian Federation (five of *Myotis dasycneme*, three of *Myotis brandtii*, five of *Myotis daubentoniid*, four of *Nyctalus noctule*, six of *Pippistrellus nathusii*, two of *Plecotus auritus* and single *Vespertilio murinus*) were analysed in the study. The products of amplificaton of RdRp genes was detected in 50% RNA samples extracted from rectal swabs of animals. Sequencing confirms the presence of merbecoviruses in three of six analysed samples from *P. nathusii* only (semi-adult animals, one of them was female and two were males). The animals were caught at the same time, in the same geographic location, so we believe they were from the same colony. BLAST against GenBank records as well as topology of the phylogenetic tree based on the fragments of the RdRp genes revealed that the bats were infected with same virus, MOW-BatCoV/15-22.

The complete genome of MOW-BatCoV 15-22 showed the highest similarity (88% identity) to MERS-related viruses from bats (Bat-CoV/H.savii/Italy/206645-40/2011 and Bat-CoV/P.khulii/Italy/206645-63/2011) which has been reported in Italy [17]. *P. nathusii* is a migratory bat, which habitats the major parts of Europe: from Fennoscandia and British Isles in the north to the Mediterranean areas in the south. The breeding areas of this species are regions of north-eastern Europe. As a result of low abundance of aerial insects during winter, Nathusius’ pipistrelles from Central European (Germany and Poland) and northeastern populations (Fennoscandia, the Baltic countries, Belarus, and Russia) perform long-distance flights migration in the late summer (during approximately two months with stopping for mating) in the Switzerland, Benelux countries, France, Spain, Italy, and Croatia [52,53]. The longest migration record of this species was documented at 2224 km, between S Latvia and N Spain [53]. During migration, *P. nathusii* may come into contact with bats of the same species when mating. And they can also come into contact with bats of other species in roosting areas. Migration routes explain the fact that very similar viruses have been found in bats from Italy and Russia.

According to phylogenetic analysis of the complete genomes the MOW-BatCoV 15-22 falls into clade of human/camel’s MERS viruses together with a few bat viruses and due to its distinct phylogenetic position and amino acid differences should be considered as a novel MERSr-CoV. The replicase polyprotein of new virus (MOW-BatCoV 15-22) showed more than 90% homology for six of seven domains, the only NSP3 (ADRP) is shown 68,7% of homology to sequences of the other members of Merbecovirus. According to demarcation criteria of ICTV [51], we believe MOW-BatCoV 15-22 represents the same species of Merbecovirus as NeoCoV because of phylogenetic analysis of RdRp showed, MOW-BatCoV 15-22 and NeoCoV together are closest to MERS-CoV. Of known bat viruses, the NeoCoV from *Neoromicia capensis* (S. Africa) is the closest to Middle East respiratory syndrome coronavirus (MERS-CoV) which could infect humans and dromedary camels and it is considered that NeoCoV shares sufficient genetic similarity in the replicase genes to be part of the same viral species with MERS-CoV [20,54].

Phylogenetic analysis of complete genome sequences as well as N- and RdRp-equences suggest that ten viruses from bats found in distinct geographic regions (MOW-BatCoV 15-22 from Russia, MG596802.1 and MG596803.1 from Italy, NeoCoV Neoromicia/5038 from South Africa, and multiple strains from bats in China - MG021452.1, MG021451.1, MG987420.1, MG987421.1, KX442565.1, KX442564.1) form a distinct phylogenetic clade with MERS-CoVs from humans and camels, with high bootstrap support. However, phylogenetic analysis of spike protein encoding genes demonstrated similarity of two of these ten bat viruses - the novel MOW-BatCoV 15-22 and NeoCoV - to CoVs from *Erinaceus* europaeus (the European hedgehog).

We believe that the proven close relationship between Spike genes of viruses from bats and hedgehogs which live in the same geographic regions (namely Europe) raises the question of the possibility of interspecies transmission in the present time. Middle East syndrome Coronavirus (MERS-CoV) likely originated in bats and passed to humans through dromedary camels. Previous work suggests that MERS-CoV originated from an ancestral virus in a bat reservoir and spilled over into dromedary camels around 40 decades ago, where it circulated endemically before emerging in humans in 2012 [6,7]. Presently camels play an important role as a constantly reservoir of MERS-CoV and transmit virus to people [55–57], while the bats are widely considered to be the evolutionary, disposable source of the virus [20]. But, besides dromedaries which are the proven source of human MERS and bats, MERS-related viruses have also been discovered in hedgehogs (Erinaceous). Hedgehog carriers of betacoronaviruses have been found in China [16], Germany [58], France [59] and Poland [60]. Phylogenetic analysis carried previously showed that betacoronaviruses from Chinese and Germanies hedgehogs (Ea-HedCoV HKU31 and BetaCoV Erinaceus/VMC/DEU/2012) were closely related to NeoCoV and BatCoV from African bats in the spike region. Therefore, authors suggested that the bat viruses arose as a result of recombination between hedgehogs and bat viruses [16]. Our independent finding of one more virus from European bat *Pipistrellus nathusii* which was found closely related to viruses from hedgehogs in the Spike region but not in N- and RdRp regions support the idea of recombination between ancestral viruses of bats and hedgehogs. Our independent finding of a novel CoV from a European bat with spike protein encoding sequences closely related to those from hedgehog MERSr-CoVs also suggests recombination between ancestral viruses of bats and hedgehogs. The overlap in geographic range between *P. nathusii* and the natural distribution of *E. europaeus* raises the possibility that this recombination represents an ancestral interspecies transmission. Our findings support the need for wider surveillance of MERSr-CoVs in both bats and hedgehogs

MERS-CoV targets a cell-surface receptor, the Dipeptidyl peptidase 4 (DPP4, also known as CD26). A receptor-binding domain (RBD) on the viral spike glycoprotein (S) mediates this interaction and is bound to the extracellular domain of human DPP4 [31,61]. The HKU4 and HKU5 merbecoviruses from Chinese bats are closely related to MERS-CoV in spike protein genes. The HKU4 viruses can use the MERS-CoV receptor DPP4, but not HKU5. Another MERS-CoV-related betacoronavirus, Hp-BatCoV HKU25 occupies a phylogenetic position between that of HKU4 and HKU5 and can binding of DPP4 protein for entry to DPP4-expressing cells, although with lower efficiency than that of MERS and HKU4 viruses [26]. At the same time, at least in some merbecoviruses (namely, Bat-CoV-PREDICT/PDF-2180 and NeoCoV) the domain RBDs of S-protein have been shown to use the ACE2 receptor, their RBDs amino acid composition and sequence differ greatly from RBD of SARS-CoV-2 (slightly more than 18% of identical aa) [62]. In the novel MERSr-CoV (MOW-BatCoV 15-22), the RBD region appears likely to interact with DPP4 across amino acids 366–624 within the S1 subunit. The amino acid composition of the RBD domain of the MOW-BatCoV/15-22 virus differs by approximately the same range from the RBD domains of those viruses that interact with DPP4 receptors (32-36,6% the same amino acids) and of viruses that interact ACE2 (33-33,7% the same amino acids). Thus, while it cannot be ruled out that MOW-BatCoV/15-22 binds to other cell receptors (e.g. ACE2), it is more likely it binds to DPP4.

Previous work demonstrated that only two mutations in the HKU4 coronavirus spike protein encoding region can make it infectious for human cells [61]. These are changes in one amino acid in two motifs each hPPC (recognized by furin proprotein convertase) and hECP (recognized by endosomal cysteine protease Cathepsin L). In MERS-CoV which causes the Middle East respiratory syndrome hPPC is Arg748-Ser749-Val750-Arg751-Ser760, hECP MERS Ala763-Phe764-Asn765 while in HKU4 hPPC is Ser746-Thr747-Phe748-Arg749-Ser750 hECP Asn762-Tyr763-Thr764. When Ser746 was changed on Arg (to make motifs recognizable by protease) and Asn762 on Ala (to destroy a potentially existing N-linked glycosylation site) fully mediates viral entry into human cells. In the MOW-BatCoV/15-22 virus hPPC motif is Pro758-His759-Ser760-Arg761 (based on MERS and HKU4). If compared with the ACE2 interacting S proteins of Human Sars-CoV-2, Bat CoV PREDICT/PDF-2180 and NeoCoV, then it is probably Ser760-Arg761-Thr762-Asn763. It is possible that substitution of either Pro758 or Asn763 the mentioned constitutions can lead to the furin cleavage site formation and it’s subsequent recognition by furin and could result in an increased ability to infect human cells.

hEPC in MOW-BatCoV/15-22 virus is Ala772-Tyr773-Pro774. Here, as in HKU4, there is no aa containing nitrogen, which removes the possibility of an N-linked glycosylation site. The second aa is Tyr773, which corresponds to the conservative aa at this site (Tyr or Phe) in coronaviruses. That is, there is probably nothing that needs to be changed. Thus, just one mutation in MOW-BatCoV/15-22 virus could well lead to infection of human cells.

Computer molecular docking modeling identified the DPP4 receptors of species that MOW-BatCoV/15-22 spike glycoprotein is likely able to bind to. All of these have overlapping habitats with people, providing opportunity for spillover in a natural setting. The lowest protein-protein binding was predicted in the interaction of MOW-BatCoV/15-22 Spike-protein and DPP4 of mouse and cat. This finding is supported by the lack of ability of MERS-CoV to infect cell lines derived from mice and cats [10,27]. The highest protein-protein binding was predicted in the interaction of MOW-BatCoV/15-22 Spike-protein and DPP4 of the bat *M. brandtii*, then of the hedgehog, *E. europaeus*. This is consistent with phylogenetic analyses and may indicate evolutionary relationships and recombination between spike protein sequences of bat and hedgehog CoVs. These, and previous reports of a wide distribution of MERSr-CoVs in hedgehogs suggest they could represent the natural reservoir of this clade of novel betacoronaviruses, subgenus Merbecovirus, for example, see [16,58–60] and that the pathway of emergence of MERS-CoV may be more complicated than currently known. Additionally, hedgehogs are increasingly kept as pets, with substantial numbers bred and shipped internationally, including in the Americas where MERSr-CoVs have not yet been reported. We suggest that hedgehogs should be considered, tentatively, as potential intermediate hosts for spillover of MERSr-CoVs between bats and humans, and that screening of captive hedgehogs should be conducted to rule out potential for future zoonotic spillover.

